# A new species in the *Anopheles gambiae* complex reveals new evolutionary relationships between vector and non-vector species

**DOI:** 10.1101/460667

**Authors:** Maite G Barron, Christophe Paupy, Nil Rahola, Ousman Akone-Ella, Marc F. Ngangue, Theodel A. Wilson-Bahun, Marco Pombi, Pierre Kengne, Carlo Costantini, Frédéric Simard, Josefa Gonzalez, Diego Ayala

## Abstract

Complexes of closely related species provide key insights about the rapid and independent evolution of adaptive traits. Here, we described and studied a presumably new species in the *Anopheles gambiae* complex, *Anopheles fontenillei*, recently discovered in the forested areas of Gabon, Central Africa. Our analysis placed the new taxon in the phylogenetic tree of the *An. gambiae* complex, revealing important introgression events with other members of the complex. In particular, we detected recent introgression with *An. gambiae* and *An. coluzzii* of genes directly involved in vectorial capacity. Moreover, genome analysis of the new species also allowed us to resolve the evolutionary history of inversion 3La. Overall, *Anopheles fontenillei* has implemented our understanding about the relationship of species within the *gambiae*complex and provides insight into the evolution of vectorial capacity traits, relevant for the successful control of malaria in Africa.

## Introduction

Species at earlier stages of speciation provide unique insights about the evolutionary forces involved in the origin of new species before demographic and selective processes blur the signals. However, the closer we are to the first signals of divergence, the harder it is to define the species concept, or even to intuit if this process will end in speciation [1]. Complex of species, closely related taxa where the species boundaries are uncertain, offers a compelling opportunity to study the “speciation continuum” [1, 2]. Unfortunately, the reproductive isolation is still incomplete to fully prevent introgression between taxa, hindering the true phylogenetic relationships [3]. On the other hand, the genetic exchange across backcrossed hybrids can favor adaptation [4]. Indeed, advantageous alleles can be selected in one species and introgressed in other favouring range expansions [5], altitudinal adaptation [6], or insecticide resistance [7], among others traits.

Most of the major malaria vectors across the world belong to species complexes with other non-vector species [8], providing a compelling opportunity to understand the rapid and independent evolution of their vectorial capacity [9, 10]. While malaria mosquitoes exhibit a wide ecological plasticity, preference for feeding on humans, and large population size, the non-vectors species display narrower geographical range, zoophilic host preference, and strong seasonal-dependence or reduced population size [11]. In Africa, three of the six major malaria vectors belong to the same complex, the *Anopheles gambiae: Anopheles gambiae, Anopheles coluzzii* and *Anopheles arabiensis* [12]. The complex is comprised of eight cryptic species [13-15], which differ in many ecological aspects, particularly in host feeding preference, breeding sites, feeding behavior and their role in malaria transmission [13, 16]. Most of the species inhabit natural habitats with none or a secondary role in malaria transmission. Indeed, adaptation to anthropogenic habitats, and therefore implementing their role in human malaria transmission, is an exception rather than the rule within the complex [16]. The *An. gambiae* complex is an example of speciation with gene flow, where species exhibit extensive genomic introgression, evidencing permeable gene flow barriers among them [3, 10], sustained by heterogenic patterns of reproductive isolation [17]. Consequently, pervasive introgression has hindered the elucidation of the correct phylogenetic relationships [18]. Besides, gene exchange between species in the complex has modulated their local adaptation capacity. For instance, the ability of *Anopheles arabiensis* to live in desiccating environments has been conferred by the introgression of the inversion 2La from *An. gambiae/An. coluzzii* [10, 19]. *Anopheles coluzzii* has developed resistance to insecticide treatments due to the introgression of the *kdr* mutation from *An. gambiae* [7]. Onwards, insecticide resistance introgressions have repeatedly occurred during the last decades [20, 21]. Thus, introgression has accelerated local adaptation and range expansion within the complex. Complex of closely related species also offer a compelling opportunity to study locally adapted alleles. Comparative genomics in recent species radiations allows unraveling the genetic basis of the traits involved in their ecological, behavioral or genetic divergence [22]. In *Anopheles,* these comparative studies has contributed to elucidate some traits involved in vectorial capacity, that in turn could be used to improve vector control strategies [9]. For instance, antennal transcriptomic comparisons between *Anopheles gambiae* and *An. quadriannulatus* have provided genomic insights on host preference evolution to humans [23]. Moreover, genome wide analysis between one fresh-water (An. *gambiae)* and one salt water (An. *melas)*species has allowed to identify regions involved into the salinity tolerance within the complex [24]. Therefore, understanding the origin and the mechanisms underlying vectorial capacity within the *Anopheles gambiae* complex is decisive for the successful control of malaria in Africa [16, 25].

During an exploratory survey at La Lope National Park (Gabon) in 2014, we discovered mosquitoes morphologically identified as *An. gambiae.* Further bioecological, behavioral, taxonomic, cytogenetic, and preliminary molecular analysis suggested the probable existence of a new taxon in the *An. gambiae* complex. Then, genome-wide phylogenetic analysis placed this potential new taxon in the phylogeny of the complex as a sister species of *Anopheles bwambae,* and in the same clade as *Anopheles quadriannulatus, An. arabiensis,* and *An. melas.* Comparative genomic analysis indicated the existence of recent introgression between the potential new species and *An. gambiae/An. coluzzii.* Genes involved were enriched for detoxification, desiccation, and olfactory perception functions, directly linked to local adaptation and host preference. These analyses also elucidated the evolutionary history of the 3La inversion within the complex that entailed multiple lost events. Overall, the discovery of a probable new taxon has evidenced the importance of new species for the understanding of evolutionary relationships of species in the *An. gambiae* complex with potential implications for a better understanding of vectorial capacity traits and ultimately malaria control.

## Results

All the specimens morphologically identified as *An. gambiae* belonged to a potential unknown taxon, within the *An. gambiae* complex, hereafter called *An. fontenillei* n.sp. This species is dedicated to our colleague Didier Fontenille and his wife Marielle, medical entomologist, who has greatly contributed to the study of mosquitoes and the development of medical entomology in Africa.

### Bio-ecology of An. fontenillei

We prospected 22 sites in the National Park of La Lope in Gabon: 17 sites in the park and 5 sites in the village of La Lope, 10-15 km away from the park sites. In total, we collected more than 1,500 mosquitoes, belonging to 13 different species. Of them, 45 adults and two larvae were morphologically identified as *An. gambiae* but presented an unexpected DNA band in the PCR assay for identification of the *Anopheles*complex species [26]. In Gabon, only three species of the gambiae complex have been recorded, and all of them can be identified based on specific PCR bands [27, 28]. The individuals of the unknown species were found in six natural sites across the park, away of any human activity or presence (Fig. 1, Table S1). All the positive sites were in the edge of forest patches and close to natural marshes frequented by wild animals (e.g. African forest buffalos and other ungulates). The presence of larvae belonging to the potential new species in two of these marshes suggested a particular affinity for sunny clay soil water collections containing turbid fresh-water of rain origin. Another *Anopheles* species, *Anopheles maculipalpis,* was collected in the same breeding site. This species is known to breed in sunny, low oxygen and generally stagnant water and it has already been found breeding in sympatry with *An. gambiae* [29]. This typology of larval habitat is very similar to that of *An. gambiae, An. coluzzii, An. arabiensis*[30], but different to the other members of the complex, such as *An. merus* or *An. melas* (mangrove swamps) or *An. bwambae* (hot thermal springs), which place the new taxon in the fresh-water group of species within the *An. gambiae* complex [16]. Although no blood-fed mosquitoes were found, we assumed a preference for feeding on animals (zoophily) due to the lack of human hosts in the sylvatic sites. Mosquitoes were sampled using BG^®^ traps baited with BG-lure a source of CO_2_ [31] and Human Landing Catches-HLC-(Fig. 1B), revealing that the potential new species can feed on humans as well. Moreover, our collections in the village of La Lope (~10-15 km away of the park sites) revealed the presence of two other members of the complex (*An. gambiae* and *An. coluzzii*). No specimen of *An. fontenillei* was found in the village (HLC and larva prospections).

**Fig.1.**
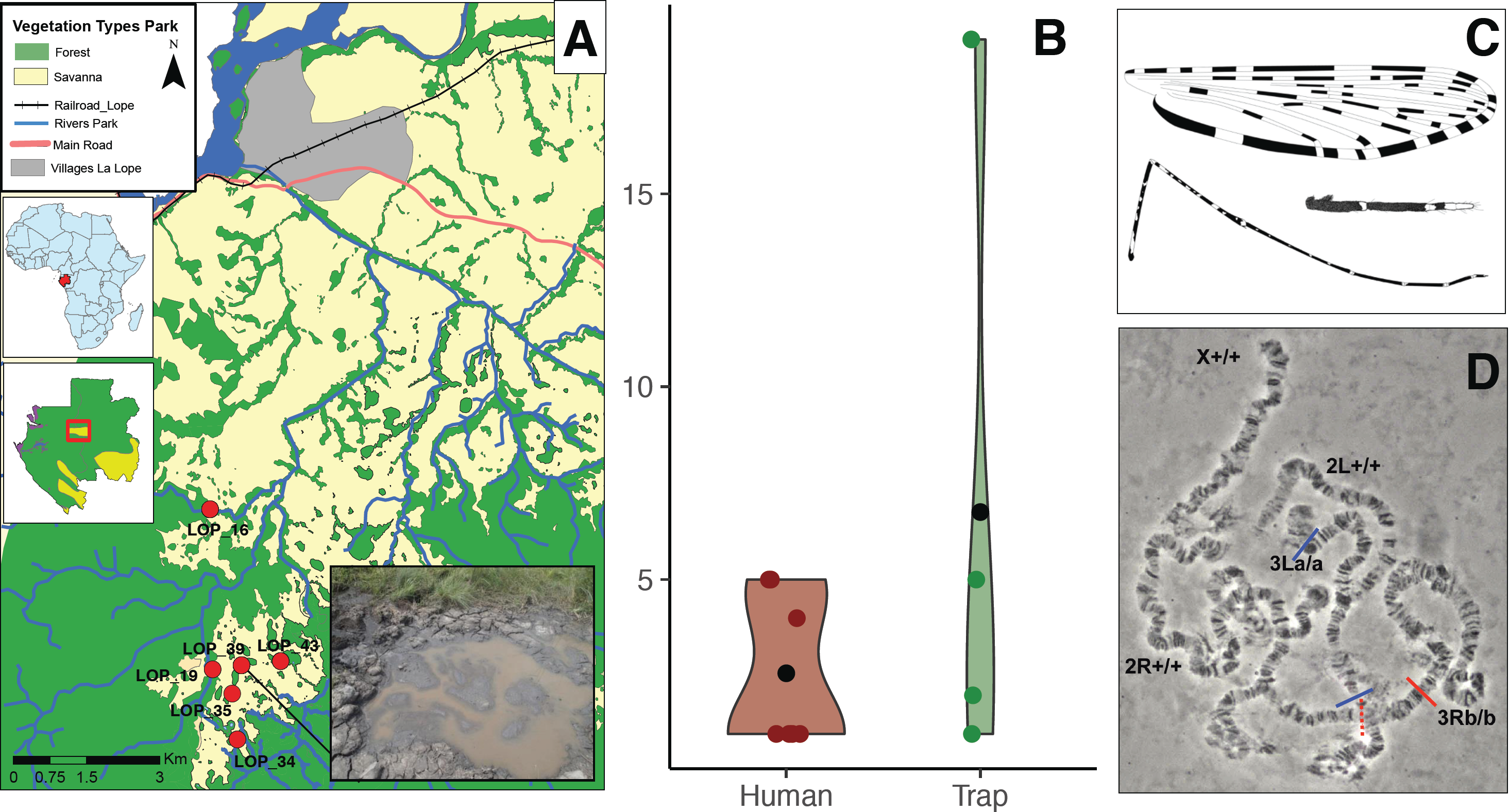
Overview of bionomic characteristics of *An. fontenillei*. (A) Geographical distribution of *An. fontenillei* within the National Park of La Lope. Breeding site where the larva of the new species was found. (B) Mean (black dots) of *An. fontenillei* collected by human landing catch (human, red) vs. BG traps (trap, green) in the park. (C) Morphological features of *An. fontenillei:* Dorsal view of the wing, maxillary palpus and hindleg with femur, tibia and tarsomeres. (D) Polytene chromosomes from ovarian nurse cells of *An. fontenillei* with a contrast-phase microscope (specimen n. 23). Chromosomal arms karyotypes are indicated following the classical nomenclature [35]. Paracentric inversions are designed by lines (red and blue) above the arms 3R(b) and 3L(a), respectively.

### Brief taxonomic description

Five *Anopheles fontenillei* specimens were preserved for taxonomic purposes (Table S1, holotypes deposited at the IRD in Montpellier, France). In general, *An*. *fontenillei*presents the classical morphotype of species within the *An gambiae* complex [15, 32, 33]: three white-scaled bands in the maxillary palpus, irregularly shaped speckling in femora and tibiae and a pale interruption on vein R1 (Fig. 1C) (for further details see Text S1). However, small differences were detected. In particular, the maxillary palpus exhibited a large white-scaled band covering completely the palpomere 5 and part of the palpomere 4 (Fig. 1C), similarly to *An. bwambae* [34].

### Cytogenetic analysis

In order to confirm the species status and its phylogenetic relationships within the gambiae complex, we collected 270 sylvatic *Anopheles* for cytogenetic purposes. Forty mosquitoes survived to attain the correct stage (half-gravid) to observe polytene chromosomes. Among them, four mosquitoes were morphologically identified as belonging to the *An. gambiae* complex, but only three revealed readable polytene chromosome preparations. According to the classical nomenclature for chromosomal rearrangements in the *An. gambiae* complex [35], all the specimens exhibited the X chromosome and the 2L arm standard arrangements, and the inversions 3Rb and 3La were fixed. In addition, the inversion 2Rl was polymorphic: inverted in one specimen and standard in the other two mosquitoes (Fig. 1D, Fig. S1). For the 2La inversion, a molecular karyotyping test is available [36]. We then used five additional specimens to validate the status of the inversion 2La [36]. All the specimens revealed a PCR-band consistent with the 2La standard arrangement, confirming our cytogenetic karyotype. Globally, *Anopheles fontenilllei* revealed a karyotype similar to *An. bwambae [34],* except for the possibility that the inversion 3Rb is fixed in the new taxon, while it is polymorphic in *An. bwambae*. Further cytogenetic works with a bigger number of individuals will be necessary to confirm the inversion polymorphisms of this species.

### Preliminary phylogenetic analysis

Sixteen specimens were used to obtain sequences for the nuclear ITS2 and IGS and the mitochondrial ND5 and COI regions, routinely used for *Anopheles* phylogenetic studies. Nevertheless, ITS2 and ND5 regions were successfully amplified and sequenced only for nine and five specimens respectively (Table S1). Overall, all the genes exhibited a low diversity with an unique haplotype, except the COI gene, which presented 5 haplotypes. The phylogenetic trees showed that *An. fontenillei* sequences always clustered with *An. bwambae* within a monophyletic clade (Figure S2), corroborating the previous cytogenetic results but in contrast with the ecological observations. Two of the four genes, ITS2 and ND5, revealed differences between *An. fontenillei* and *An. bwambae* (Figure S2). These results are in congruence with previous studies revealing that most of the classical molecular markers are not discriminant among species in the complex due to their extensive introgression [10].

Overall, the new taxon revealed important similarities to *An. bwambae*, a thermal spring breeding species from a forested area of Uganda (Semliki valley). Taxonomic (large band in the palpomeres 4 and 5), cytogenetic (chromosomal inversions) and molecular (sequence divergence) criteria could not differentiate between the two species. On the other hand, ecological (freshwater marshes vs thermal springs) and geographical (allopatric distribution: Gabon vs Uganda) results clearly discriminated between *An. fontenillei* and *An. bwambae*. Therefore, further genomics studies are needed to elucidate the true phylogenetic place of *An. fontenillei* within the *An. gambiae* complex.

### Anopheles fontenillei is a potential new species of the Anopheles gambiae complex

We conducted a genome-wide analysis in order to accurately locate the new species in the *An. gambiae* complex phylogenetic tree. According to previous studies [10], we considered that the true *An. gambiae* complex species tree is mainly observed in the X chromosome. Hence, we initially focused on analyzing the X chromosome arm. For this purpose, we made a genome assembly of one *An. fontenillei* individual sequenced at high coverage (~112X) (Table S2). This assembly was nearly complete, according to the 96% of completely found BUSCO genes, but highly fragmented with a N50 of 21kb (Materials and methods, Table S3B and Table S3C). We then added to the available multiple alignment file (MAF), based on six described gambiae complex species [10], our *An. fontenillei* assembly and the highest coverage *An. bwambae*individual publicly available (see Material and Methods). Maximum likelihood (ML) phylogenetic trees were built for each non-overlapping 50kb windows (see Materials and Methods, Table S4).

Following this approach, the relationship among species observed in the X chromosome was as shown in Figure 2. Similarly to previous studies, the relative position of the basal node, *An. merus* and *An. coluzzii-An. gambiae* clade, was not clearly determined due to incomplete linage sorting (ILS, [10]. In our analysis of the X chromosome, *An. fontenillei* appeared as the sister species of *An. bwambae* in 83% of the trees (264 out of 319). However, there exists a certain ambiguity to determine the ancestral taxon of the clade. While, the clade branches with *An. quadriannulatus*in 78 of 319 windows (Figure 2), it branches with *An. arabiensis* in 59 of 319 windows (Fig. S3). Assuming that the X chromosome shows the true species tree, we then presume that either *An. quadriannulatus* or *An. arabiensis,* shared a common ancestor with *An. fontenillei* and *An. bwambae.*

**Fig.2.**
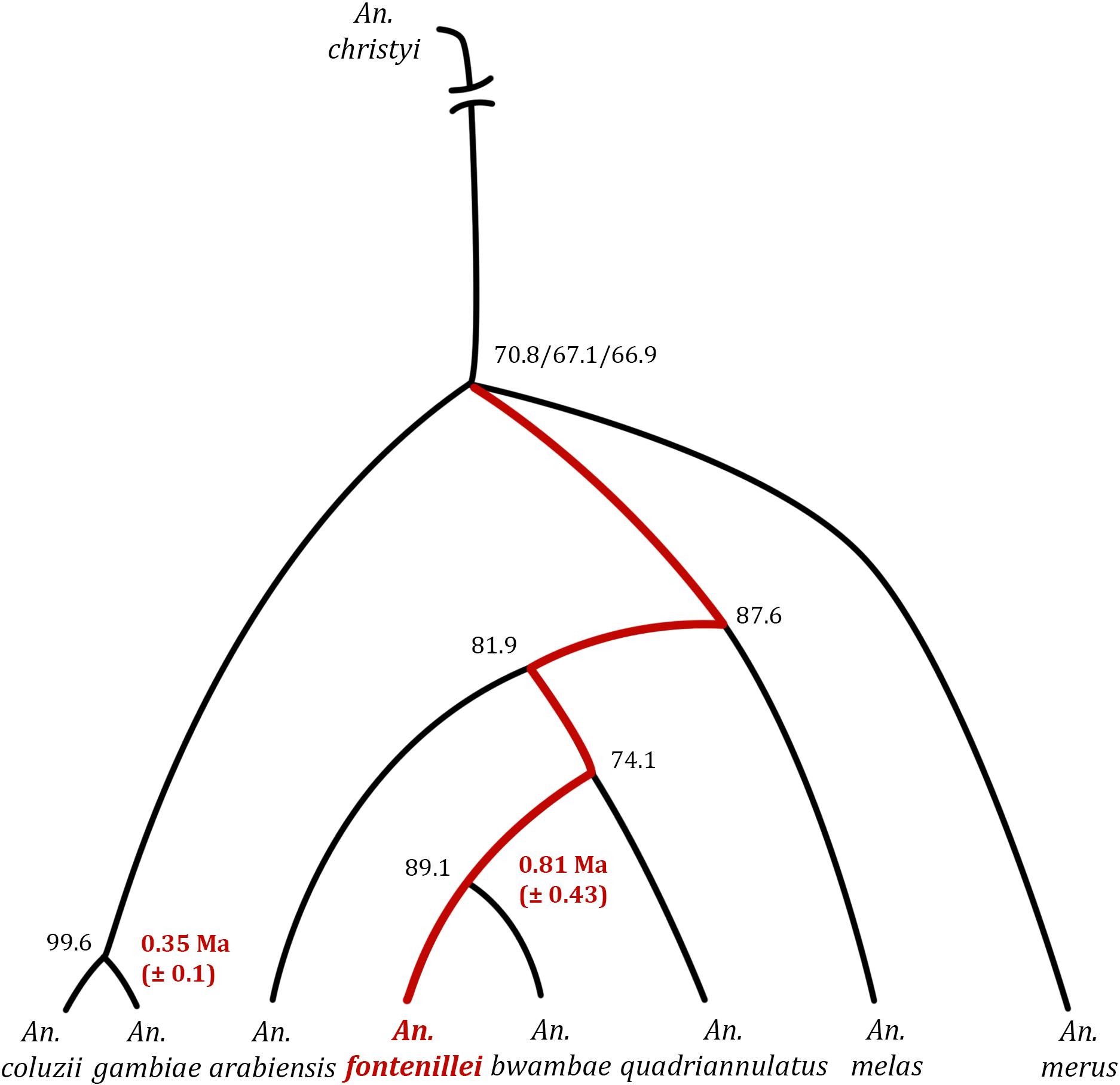
Most common species tree. 78 windows in the X chromosome show this tree topology with a week disagreement in the basal node. Black numbers represent bootstrapping values and in red the millions years estimated based on the pairwise distances of the ML phylogeny and assuming a substitution rate of 11.x10^-9^ per site, per generation and 10 generation per year.

To investigate if we find a stable distinction between *An. fontenillei* and *An. bwambae,* we repeated the analysis creating a new MAF adding 3 additional individuals of *An. fontenillei,* and 2 additional individuals of *An. bwambae* (see Material and Methods). Out of the 343 analyzed windows, 278 (81%) showed trees where individuals of *An. fontenillei* and *An. bwambae* clustered together and these two species were permanently separated, indicating that they are different populations and/or species (Fig. S4).

We also estimated the pairwise genetic distance between *An. fontenillei* and *An. bwambae* and compared it with the pairwise genetic distance between *An. coluzzii* and *An. gambiae,* the most recently diverged species within the complex (Fig. S5A) [10, 37]. The pairwise genetic distance was significantly larger in the *An. fontenillei*-*An. bwambae* clade compared with the *An. gambiae*-*An. coluzzii* clade (bootstrapping analysis, median 0.0117 and 0.0067 respectively, Figure S5B, Table S5). If we assumed a substitution rate of 1.1x10^-9^ per site, per generation, and 10 generation per year [38], there had been 0.53 Ma since the *An. fontenillei*-*An. bwambae* clade split, and 0.31 Ma since the *An. gambiae*-*An. coluzzii* clade split (Fig. 2 and Figure S3).

This result together with the clear ecological distinction between *An. fontenillei* and its closest species within the complex, *An. bwambae,* suggested that *An. fontenillei* is a new species in the *An. gambiae* complex rather than a sub-population of the *An. bwambae* species.

### Recent and ancestral relationship of An. fontenillei with other members of the complex

We extended our analysis from the X chromosome to the whole genome. In 84% of the analyzed genome, *An. bwambae* is the closest species to *An. fontenillei,* forming the *An fontenillei-An. bwambae* (FB) clade (Fig. 3, R line, Figure S6, Table S5). This proportion is similar in every chromosome arm ranging from 78.4% in the 3R chromosome arm to the 86.6% in the 3L chromosome arm. The proportion of FB clade in the autosomes, 84.1%, is in concordance with the FB clade proportion in the X chromosome, 82.8%, *i.e* the species tree, indicating that *An. fontenillei* has not extensively introgressed with other members of the complex in a recent period. However, the relationship of the FB clade with its closest species or other clades, showed a very different pattern between the X chromosome and the autosomes (Figure 3, A line). In the X chromosome, the majority of windows showed the species tree, as it was previously described [10]. Accordingly, FB clade is closely related to *An. quadriannulatus* (27.5%) or *An. arabiensis* (24.5%). While the autosomes, the majority of windows showed the recent introgression between *An. arabiensis* and *An. gambiae*-*An. coluzzii* clade, the A(GC) clade[10]. In the autosomes, the FB clade is branching with the A(GC) clade for the majority of windows, 27.6% (Fig. 3, Figure S6, Table S6). The next more frequent topology, 16.4%, shows the FB clade with *An. quadriannulatus* as the closest species. However, if we do not take into account the 2La inversions (see below), which shows its own topology, this proportion was reduced to 9.6%. According to the X chromosome analysis, we could not conclude whether *An. quadriannulatus* or *An. arabiensis* is the closest species to the FB clade due to similar number of windows showing one or the other topology (Fig. 2, Fig. S3). However, in the autosomes we can clearly observed that the FB clade is more frequently branching with A(GC) clade. Hence, if *An. quadriannulatus* is the closest species to the FB clade, the FB common ancestor must have suffered introgressions with *An. arabiensis* prior to the *An. arabiensis* and *An. gambiae-An. coluzzii*introgressions. On the contrary, if *An. arabiensis* is the closest species to the FB clade, there is no need of additional introgression to explain the observed results. Only that, the split of the FB common ancestor with *An. arabiensis* must have been prior to the introgressions between *An. arabiensis* and the GC clade.

**Fig.3.**
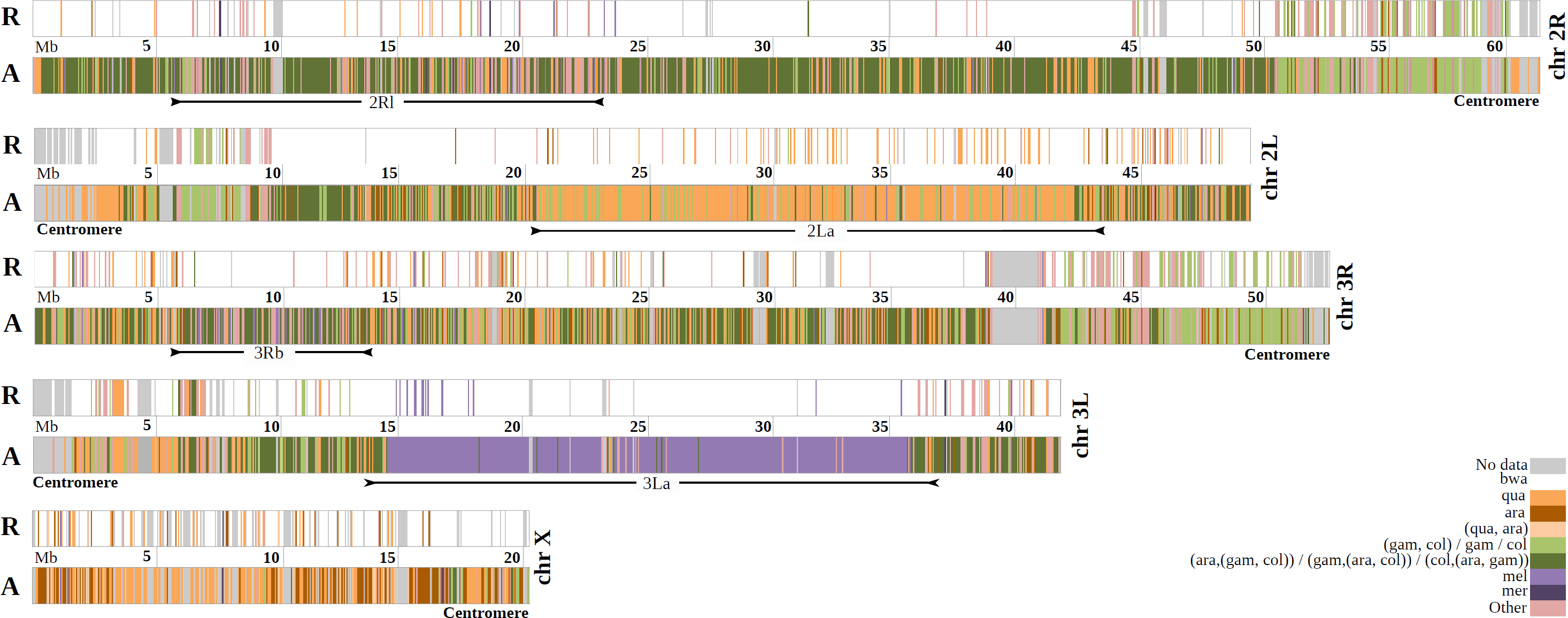
Recent (R) and ancestral (A) relationship of *An. fontenillei* **with other species in the *An.* gambiae complex according to the phylogenetic trees in 50kb non-overlapping windows along each chromosome arm.** (R) *An. fontenillei* closer species or clade on each tree. (A) When the closer species in the tree is *An. bwambae,*then *An. fontenillei*-*An. bwambae* clade closer species or closest clade is shown for each window.

Most of the windows that do not show the FB clade are located close to the centromeric ends. This is mainly observed, in the 2R and 3R last ~11Mb close to the centromere and, a similar pattern is observed in the first ~10Mb close to the centromere of the 2L chromosome arm (Fig. 3, R line). In these regions, the proportion of windows showing the FB clades is smaller than in the rest of the chromosome. Interestingly, another difference is that the proportion of trees showing *An. fontenillei* close to GC clade or either *An. gambiae* or *An. coluzzii* is substantially bigger than in the rest of the chromosome. Specifically, in regions close to the centromeres, the FB clade proportion is ~40% for 2R and 3R, and, 51% for 2L chromosome arm while in the rest of the three chromosome arms, the FB clade proportion is > 80%. The proportion of trees showing *An. fontenillei* close to GC clade or *An. gambiae* or *An. coluzzi* is ~20% for 2R and 3R, and, 7% for 2L chromosome arm while on the rest of the three chromosome arms this proportion is < 1%. We checked that the alignment quality of these regions were not different from other regions in the chromosome to exclude possible biases due to low quality alignments (Fig. S7, Materials and Method). The alignments in these regions are shorter but still have on average 16,482 informative positions per window and they showed better alignment qualities than in the other regions of the chromosome arm, in terms of the proportion of gaps or alignment fragmentation. Hence, neither low quality nor short alignments are likely to be the cause of the observed differences (Fig. S7, Materials and Methods).

Although we cannot discard that these regions are a consequence of incomplete lineage sorting, it is difficult to explain why the FB clade appears close to the GC clade repeatedly. If we remove the FB clade from the analysis, we cannot observe any differences in these regions compared to the rest of the genome. We argue that these windows may indicate a very recent introgression between *An. fontenillei* and *An. gambiae* or *An. coluzzii*, or both.

### Recent introgressed genes are enriched in metabolic detoxification, desiccation, and olfactory perception

We analyzed the gene content of windows were *An. fontenillei* instead of branching with its closer species, *An. bwambae,* clustered with the major malaria vectors i) *An. gambiae*, ii) *An. coluzzii*, or iii) the GC clade. These species occur in sympatry at La Lope area, so it could be possible that they share DNA through secondary contact. We analyzed the three ML tree topologies related with this possible recent introgression separately because the presence of the 2La polymorphic inversion may affect the results: the inversion breaks apart the more frequently observed GC clade, because the *An. coluzzii* individuals used for this study were predominantly inversely oriented while *An. gambiae* individuals were predominantly standardly oriented.

i) There were 64 windows harboring 198 genes were *An. fontenillei* branched with *An. gambiae*. We performed a functional enrichment analysis of the 198 genes, with DAVID, using *An. gambiae* genome as background [39, 40]. There were four significant clusters (Table 1). The first three clusters were related with cuticle proteins, membrane transporter activity, peptidases and proteases. All these protein families had been related to metabolic detoxification of insecticides in high-throughput genome-wide studies in several mosquito species (reviewed in [41]). Additionally, the cuticle proteins had also been described as being critical for the desiccation tolerance in embryos [42]. Interestingly, the GO term of the peptidases and proteases cluster had been previously related with high evolutionary rates [9]. The last cluster, was related to *heat shock protein 70*, a conserved protein related to heat stress but also to oxidative stress and detoxification of some toxines [43, 44]. The InterPro domain in this cluster has also been shown to be a rapid evolving gene family (Table1, [9]).

**Table 1.** Functional enrichment analysis result. Significantly enriched clusters (>1.3 enrichment score) obtained with the analysis of functional terms with DAVID.

| Tree topology | # of windows | # of genes | Term summary | Enrichment Score | GO terms | InterPro domains |
| --- | --- | --- | --- | --- | --- | --- |
| (F,G) | 64 | 198 | Insect cuticle protein | 5.6 | GO:0042302 | IPR000618 |
|  |  |  | Transmembrane transporter activity | 1.6 | GO:0042391, GO:0015701, GO:0015301, GO:0019531, GO:0015106, GO:0015116, GO:0008271, GO:0051453, GO:0005254, GO:1902476, GO:0005887 | IPR001902, IPR011547, IPR002645 |
|  |  |  | Peptidase activity | 1.6 | GO:0004252 | IPR001254, IPR018114, IPR009003 |
|  |  |  | Heat shock p70 | 1.4 | - | IPR018181, IPR013126 |
| (F,C) | 25 | 62 | None | <1.3 | - | - |
| (F,(GC)) | 35 | 89 | Flavin monooxygenase | 3.6 | GO:0004499, GO:0050661, GO:0055114, GO:0050660, GO:0004497 | IPR000960, IPR020946, IPR023753 |
|  |  |  | Olfactory receptor | 1.3 | GO:0050911, GO:0004984, GO:0005549, GO:0005886 | IPR004117 |
| (F,L) | 15 | 25 | Larval midgut histolysis | 4.9 | GO:0035069, GO:0097200, GO:0097194, GO:0005737 | IPR002138, IPR001309, IPR015917 |

ii) There were 25 windows containing 62 genes where *An. fontenillei* clusters with *An. coluzzii*. In these case, none of the clusters were significantly enriched for a particular functional term. Finally, iii) there were 35 autosomal windows containing 89 genes in which *An. fontenillei* branched with the GC clade. If these windows are actually regions of recent introgression, this would mean that those genes were introgressed between *An. gambiae* and *An. coluzzii* common ancestor with *An. fontenillei*. The functional term enrichment analysis, showed two significantly enriched clusters (Table 1). The most significant cluster was enriched in Flavin monooxygenase, which share function similarity with the cytochrome P450-monooxygenases [45]. The P450 proteins are one of the main protein families related to metabolic detoxification of insecticides in mosquito species (reviewed in [41]). The other significant cluster is related to olfaction. Three of the four GO terms in the cluster (GO:0050911, GO:0004984 and GO:0005549) and the InterPro domain (Table 1) had been described to show high evolutionary rates [9].

Following a reverse complementary approach, we also checked whether known mutations that confer resistance to insecticides or to some infections, both traits relevant for malaria transmission, where present in *An. fontenillei.* Specifically, we checked 42 mutations in 14 genes related to insecticide resistance [20] and five mutations in one gene related to immunity and infection resistance [46]. We map the four *An. fontenillei* individuals to the reference genome (AgamP3) with *bwa-mem*(Text S2.4). Only two mutations in two genes related with insecticide resistance were found in this analysis (Table S7). GSTE6 mutation (E89D) and GSTE3 mutation (N73I) were observed in the four sequenced *An. fontenillei* individuals. The GST protein family, together with the P450 family, are considered to be determinant in the metabolic detoxification of insecticides in mosquitos (reviewed [41]). We checked whether this mutation was present in the other members of the complex. All the available genome references, *An. gambiae* Pimperena, *An. coluzzii, An. quadriannulatus*, and *An. arabiensis,* showed the susceptible mutation. However, in the MAF made with wild specimens, all the species that could map to those regions, *An. gambiae, An. coluzzii* and the three *An. bwambae* individuals showed the resistant alleles, as did the four *An. fontenillei* individuals. This showed that this mutation is polymorphic in several species within the complex and suggest that the resistant phenotype should thus also be shared by all these species.

### Chromosome inversions reveal putative introgression events in the Anopheles gambiae complex

There are two main inversions in the *An.* gambiae complex, which emerged in our phylogenetic analysis, and shaped the chromosomal evolution within the complex: the 2La inversion and the 3La inversion. The 2La inversion has only been described in *An. arabiensis, An. gambiae,* and *An. coluzzii [35].* Neither *An. bwambae* nor *An. fontenillei* have this inversion. Hence, in this region of the 2L chromosome arm the FB clade is closer to *An. quadriannulatus,* defining a well-determined different block easily distinguishable in Fig. 3 (line A). On the contrary, *An. fontenillei* had the 3La inversion fixed as well as *An. bwambae* and *An. melas,* as shown by the cytogenetic results. In the 3L chromosome arm the inverted region can also be easily identified because in those windows, the FB clade is closely related to *An. melas* (Figure 3, line A). The inferred breakpoints based on the ML tree topology of the 2La and 3La are inside the known cytological breakpoint ranges, except for the 2L telomeric breakpoint, which was 400kb shorter (Table S8, [35], VectorBase.org).

The majority of windows, 45%, in the 3L chromosome arm showed the three known *Anopheles gambiae* complex species with the 3La inversion, *An. fontenillei*, *An. bwambae* and *An. melas*, together and separated from the species without the inversion: *An. arabiensis*, *An. quadriannulatus*, *An. merus*, *An. gambiae*, and, *An. coluzzii* (Figure 4, Figure S6). Additionally, this topology also suggests two events of introgression, i) *An. arabiensis* with the *An. gambiae* and *An. coluzzii* common ancestor and ii) *An. merus* with *An. quadriannulatus* (Figure 4). To date the 3La inversion, we estimated the pairwise distances between *An. fontenillei* and *An. quadriannulatus* in the 3L chromosome arm outside and inside the inversion (3L: 14.5-35.9 Mb+/-the 500Kb flanking region). Outside the inversion, the divergence between *An. fontenillei* and *An. quadriannulatus* was 1.4 Ma (+/-0.91), which is similar to the one estimated in the more common X chromosome phylogenetic tree between *An. fontenillei* and *An. quadriannulatus* (1.25 Ma (+/-0.54), Figure S8, Table S9). We then estimated the divergence between *An. bwambae* and *An. quadriannulatus* outside the inversion and again it was similar to the one previously estimated for the more common X chromosome phylogenetic tree between these two species: 1.24 Ma (+/-0.6). However, the divergence estimated inside the inversion between *An. fontenillei* and *An. quadriannulatus*, and *An. bwambae* and *An. quadriannulatus* were 2.53Ma (+/-0.97), and 2.23 Ma (+/-0.76), respectively. These estimates are on the range of the *Anopheles gambiae* complex origin around 2 (+/-0.64) Ma ago. We repeated this analysis using *An. arabiensis* instead of *An. quadriannulatus* and we obtained similar results (Table S9). We could not accurately date the 3La inversion with this method due to the high uncertainty, but we could show that the inversion is at least older than *An. melas*, *An. arabiensis*, *An. quadriannulatus*, *An. bwambae*, and *An. fontenillei* group. However, *An. arabiensis*and *An. quadriannulatus* showed the standard karyotype of the 3La inversion. According to the phylogenetic trees, we thus hypothesized that the ancestral karyotype of the group is the 3La inversion and that *An. quadriannulatus* lost the inversion in the introgression from *An. merus*, as has already been suggested by

**Fig.4.**
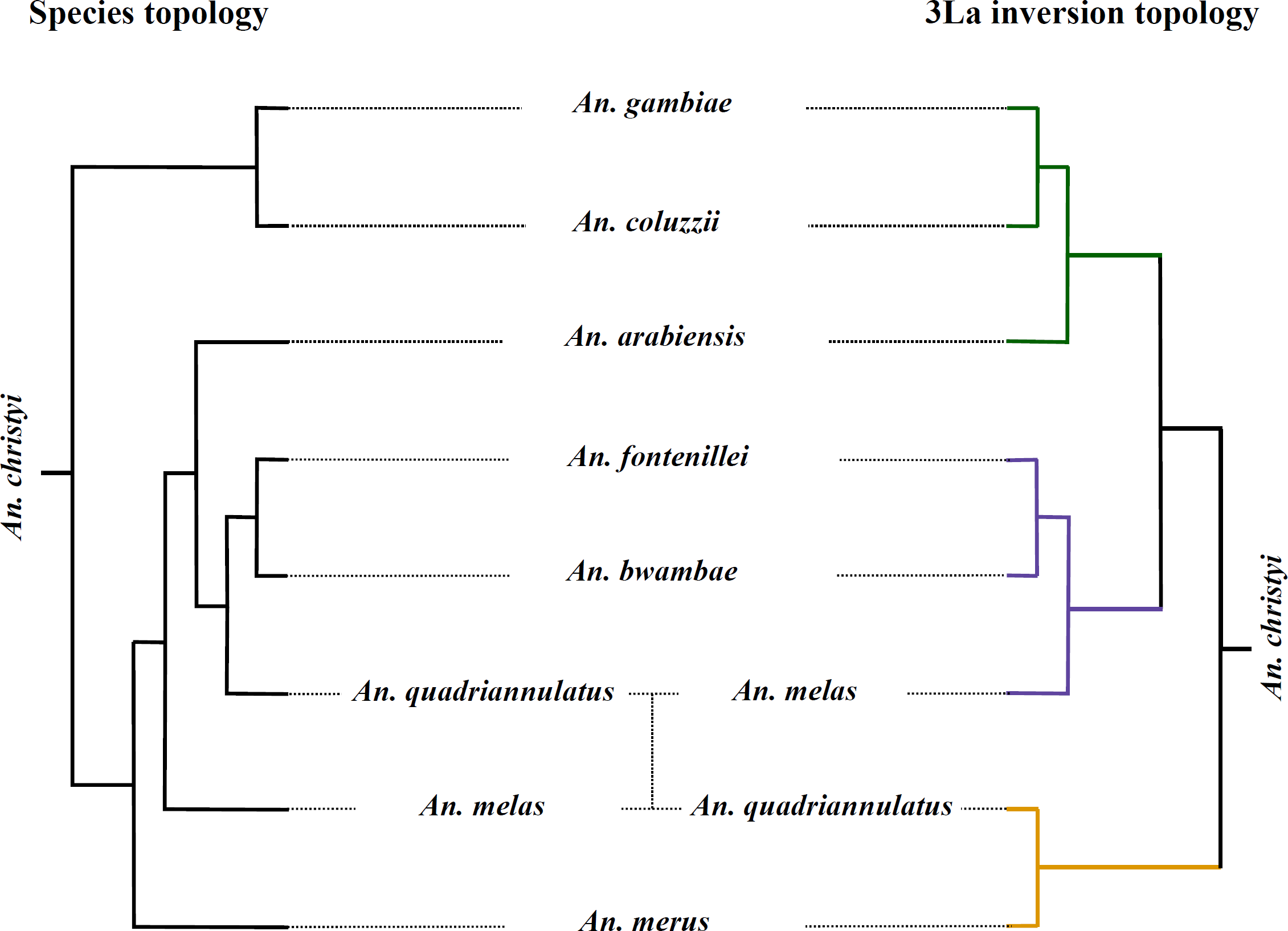
Species topology estimated from the X chromosome compared with the topology of the 3La inversion. *An. christyi* was used as outgroup species. Green color: *An. arabiensis*-GC common ancestor possible introgression. Purple color: species that share the inversion. Yellow color: *An. quadriannulatus* and *An. merus*possible introgression event.

Fontaine et al. [10], and that *An. arabiensis* lost the inversion in the introgression with

### An. gambiae/An. coluzzii

We found some interesting windows in the 3La inversion were *An. fontenillei* was closer related to *An. melas* than to *An. bwambae* (Figure 3, line R). In this case, these windows may be related with regions nearby the inversion breakpoints maintained through positive selection. We made a functional enrichment analysis of these windows using DAVID. There are 15 windows containing 25 genes that showed this topology. There was only one enriched cluster with genes related to the stage-specific breakdown of the larval midgut during metamorphosis, that allow replacement of larval structures by tissues and structures that form the adult (Table 1).

Finally, the 3Rb inversion and the 2Rl polymorphic inversions revealed by the karyotyping of *An. fontenillei* individuals do not leave a clear pattern in the genomic analysis performed here (see Fig. 3). Both inversions are only shared with *An. bwambae,* the closest species to *An. fontenillei,* and hence it is not expected to observed big differences in those regions.

## Discussion

In 1975, the English entomologist G. B. White wrote: “As time passes, it becomes increasingly less likely that other sibling species of this complex (An. *gambiae)* will be found” [47]. Indeed, during the last 40 years, only one new species, *An. quadriannulatus B* (recently called *An. amharicus),* has been discovered [15, 48], and *An. coluzzii* has been separated from its sister species, *An. gambiae* [15]. In 2014, we discovered a potential new species belonging to the *An. gambiae* species complex. The species, *An. fontenillei,* was found in a mosaic savanna-forest area of Gabon, Central Africa. This region is characterized by hosting the last vestige of savanna in the Congo rainforest basin [49]. However, this habitat is not unique, and other parts of Gabon and Central Africa could entertain the presence of this species. The new mosquito seems to breed in rain dependent, sunlit, and open pools, evidencing similar larval ecology to other fresh-water species within the complex [16]. According to its ecology, we presumed a zoophilic host preference (Fig 1B). This behavior has already been found in other members of the complex, such as *An. quadriannulatus* [50], and it seems an ancestral character. However, *An. fontenillei* can also feeds on humans, therefore, showing a generalist feeding habit with potential consequences on parasite transfer between human and animals [51]. Indeed, the ancient and recent history of La Lope provided multiple opportunities for *An. fontenillei* to adapt to humans [52]. In the Neolithic age, La Lope was commonly colonized for hunting by nomad tribes, and in the last century there was a forestry industry in the park. However, whether this trait is ancestral or recently acquired (i.e. by introgression, see below) will need further investigations (Table 1).

In order to disentangle its phylogenetic position within the complex, we sequenced and *de novo* assembled *An. fontenillei* genome. The new genome allowed us to determine that *An. fontenillei* and *An. bwambae* are sister species. Pairwise comparisons revealed a higher divergence time between *An. fontenillei* and *An. bwambae* than between *An. gambiae* and *An. coluzzii* (Fig. 2, [37]), corroborating the geographical and ecological assumptions of two different species (Fig. 1). The *An. fontenillei*-*An. bwambae* (FB) clade was placed together with *An. quadriannulatus*, *An. arabiensis* and *An. melas,* being *An. quadriannulatus* or *An. arabiensis* the closest species of the clade (Fig. 2). This is, to date, the most exhaustive phylogenetic tree of the complex, including eight of the nine species described (no genome sequence is available for *An. amharicus).*

Consistent with Fontaine *et al.,* [10], we found pervasive evidence of introgression in *An. fontenillei,* confirming the permeable species boundaries in the *An. gambiae*complex [37, 53]. Introgression within species complexes is common in nature, challenging the possibility to trace the evolutionary history of species [3]. Interestingly, we observed patterns of recent introgression between *An. fontenillei* and the clade *An. gambiae-An. coluzzzi* (GC), particularly in the centromeric regions (20% of the phylogenetic trees). These last two species were found in the village close to the sylvatic sites where *An. fontenillei* was sampled (La Lope, Fig. 1A), indicating a potential contact among them. The genomic windows introgressed were mostly enriched for genes associated with detoxification, desiccation tolerance, and olfactory perception (Table 1), which have been related with enhanced vectorial capacity [9]. These traits allow species to inhabit a broader range of habitats, and blood-feeding on different hosts. The evidence of recent gene exchange between *An. gambiae-An. coluzzii* with other species of the complex, may alter the evolution of these two major malaria vectors, with potential consequences for malaria transmission *(i.e.* adaptation to sylvatic habitats and/or preference for feeding on animals). However, we cannot discard that this patterns of recent introgression in centromeric regions could be affected by the low recombination rate in those areas, that could help to protect introgressed haplotypes for a longer time compared with other genomic regions [54]. Finally, we resolved the evolution of the inversion 3La in the *An. gambiae* complex While, this inversion was thought to be present in the ancestor of *An. melas* and *An. bwambae*, we estimated that the origin of this inversion predated the radiation of the gambiae complex [34]. Moreover, we evidenced that the inversion was independently lost by *An. arabiensis* and *An. quadriannulatus* (Fig 3, Fig. 4). Although the 3La inversion has not been associated yet to any trait, we observed functional enrichment in larval midgut histolysis genes in recently introgressed regions between *An. melas*and *An. fontenillei* (Table 1). Again, these two species are present in Gabon, and could envision potential gene exchange between them. Chromosomal rearrangements have modulated the evolution of multiple species by affecting local adaptation or speciation [5, 55-60]. In our genomic analysis (Fig. 3), we also observed the genomic signature of the 2La inversion that affects the phylogenetic relationship between *An. fontenillei*, *An. arabiensis*, and *An. quadriannulatus*, highlighting the impact of fixed inversions in chromosome evolution within the complex.

Besides the titanic collection effort led in Africa during the last century, the rainforest of Central Africa has carefully hidden a new piece in the jigsaw puzzle of the *An. gambiae* species complex. The discovery of a new species in the *An. gambiae*complex has provided new insights into genome evolution (i.e. inversion 3La) and local adaptation (i.e. salinity tolerance) in this group of closely related species. Moreover, the new species has been an active actor in the evolution of *An. gambiae-An. coluzzii*, exchanging genes involved in vectorial capacity. These introgressions open new questions about how local populations of the major vectors, *An. gambiae and An. coluzzii*, have been affected. Indeed, adaptation to rainforest habitats, host preference or resting behavior could have been modified at La Lope. New studies may provide important insights about how vectorial traits have evolved from wild to domestic populations within the complex, with a direct impact in future malaria control strategies.

## Material and Methods

### Research and ethics statements

We sampled *Anopheles* specimens under the national park entry authorization AE16008/PR/ANPN/SE/CS/AEPN and the national research authorization AR0013/16/MESRS/CENAREST/CG/CST/CSAR. Moreover, we obtained the approval by National Research Ethics Committee of Gabon (0031/2014/SG/CNE) to perform the human-landing catch (HLC) collections.

### Mosquito sampling and species identification

Mosquitoes were sampled in the National Park of La Lopé in Gabon, Central Africa, in an exploratory survey in November 2014. Since, several collections were carried out in June 2015, February 2016 and November 2016. (Fig. 1, Table S1). Adults were collected using BG traps with BG-lure and a source of CO_2_ and HLC, while larvae were sampled by the dipping method [61]. Collected *Anopheles* mosquitoes were taxonomically identified according to standard morphological features [32, 33]. Then, they were individually stored in 1.5 mL tubes at-20°C and sent to CIRMF for molecular analysis. Total genomic DNA from specimens morphologically identified as belonging to the *An. gambiae* complex was extracted using the DNeasy Blood and Tissue Kit (Qiagen) according to the manufacturer’s instructions. Genomic DNA was eluted in 100 μL of TE buffer. A first molecular diagnostic (PCR-based) was performed to molecularly identify species within the complex [26]. Surprisingly, an unspecific fragment of 700 bp was amplified. This band does not correspond to any of the species reported by the PCR-RFLP diagnostic test [26].

### Mosquito karyotyping

Half-gravid females were sampled in November 2016 (Table S1) in forest sites where we previously found the unspecified taxon. Females were collected by HLC and feed to complete their blood-meal. Mosquitoes were allowed to develop follicles for 25 h at field temperature. Then, ovaries were dissected and stored in Carnoy’s fixative solution (three parts 100% ethanol: one part glacial acetic acid, by volume). At the CIRMF, we squashed the ovaries in a drop of 50% of propionic acid to obtain the polytene chromosomes [62]. The banding patterns of polytene chromosomes were examined using a Leica DM2000 and a camera system Leica DFC 450 (Leica Microsystems GmbH, Wetzlar, Germany). Chromosomal arms and inversions were recorded and scored according to *An. gambiae* chromosome map [63].

### Preliminary sequencing analysis

In order to obtain further information about the unrecognized PCR band, we sequenced three genes previously employed for phylogenetic studies in the complex following the authors’ instructions: internal transcribed spacer subunit 2 (ITS2~490 bp [64]); NADH dehydrogenase subunit 5 (ND5~300 bp [65, 66]); and cytochrome c oxidase subunit I (COI~495 bp [67]). Moreover, we designed a new set of primers for amplifying a fragment of the intergenic spacer gene (IGS~267 bp; IGSKPF 5’-CTCTTGTGAGAGCAAGAGTGT-3 ’ and IGSKPR 5’-ATCAAGACAATCAAGTCGAGA-3’) used also for species identification in the complex. For the IGS gene, PCR reactions were carried out in 25μl reaction volume than included 1X Qiagen PCR buffer (Qiagen, France), 1.5mM MgCl2, 200μM each dNTP (Eurogentec, Belgium), 10 pmol of each primer, 2.5 U Taq DNA polymerase (Qiagen, France) and 1-20 ng of template DNA. Amplifications were performed using a Mastercycler Gradient thermocycler (Eppendorf) under the following conditions: an initial step at 94°C for 5 minutes is followed by 35 cycles of 30 seconds at 94°C, 30 seconds at 54°C, 1 minute at 72°C and a final elongation step of 10 minutes at 72°C. Five microliters of the PCR product were analyzed by electrophoresis on 1.5% agarose gels containing 0.5 μl/ml ethidium bromide and photographed under UV light.

The sequences obtained for the four regions were analyzed using *Geneious* R10 [68]. We aligned the consensus sequences for each gene with randomly chosen sequences of each species within the complex. We selected unique haplotypes to be included in the phylogenetic analysis. The best substitution model for each gene was identified using SMS [69]. The phylogenetic trees were then performed by maximum likelihood (ML) method using PhyML [70], with nearest neighbour interchange (NNI) for tree searching and approximate likelihood-ratio test (aLRT SH-like, [71]) for branch support. Visualisation of trees was done using iTOL v.3.4.3 [72].

### Genome Sequencing and Assembly

We sequenced four individuals of the unknown species using the Illumina platform at the CNAG (Barcelona). To make a *de novo* genome assembly of this species one of the individuals was deeply sequenced to ~112X. The other three individuals were sequenced at an average coverage of ~29X. All reads were paired-end 126 bp long (Table S2).

The genome assembly of the more deeply sequenced *An. fontenillei* individual was performed at the Bioinformatics Unit, CRG (Barcelona) (Table S2, S3A). Reads were trimmed and filtered using Skewer version 0.2.2 [73] to remove the adapter sequence and trimming the low quality part. A FastQC analysis was performed to check the quality of the trimmed reads. We looked at the presence of contaminants in a Kraken database, which includes complete bacterial, archaeal, and viral genomes in RefSeq [74]. We only found an enterobacteria phage phiX as contaminant (Table S3B). Then, we assembled the trimmed reads by using Platanus software version 1.2.4 [75] producing contigs and scaffolds using the paired-end information. To join the contigs within the same scaffolds, stretches of N need to be added. To fill those gaps we used Platanus *gapclose* function using the original reads (Table S3 A). At this point, improve the scaffolding of the assembled genome by using the proteins described for AgamP4 reference in VectorBase (www.vectorbase.org). We used Blat [76] to map the proteins to the assembled scaffolds and reorder and join scaffolds accordingly with PEP_scaffolder [77]. We made another round of gap filling, and due to format incompatibilities, this time we used *GapCloser* tool from the SOAPdenovo package [78] (Table S3A). To evaluate the quality of our assembly, we scanned for the presence of conserved genes among the diptera order by using BUSCO software [79]. We used 2,799 gene models conserved among diptera, and classify those genes as: i) completely found in a single sequence, ii) fragmented in different sequences, or iii) completely missing. Most of the BUSCO genes, 96%, were completely found in a single sequence (Table S3C). We finally performed a polishing step by removing the scaffolds mapping to previously found contaminants (Table S3A).

### Phylogenetic analysis

To make the genome-wide phylogenetic tree by window analysis, we took advantage of the available multiple alignment file (MAF) for six species of the *An. gambiae*complex including two outgroup species: *An. christyi* and *An. epiroticus* [10]. Briefly, we used the alignment formed by whole genome sequences from population samples of multiple individuals of *An. gambiae*, *An. coluzzii*, *An. merus*, *An. melas*, *An. quadriannulatus* and *An. arabiensis*. The *An. gambiae* PEST v3 (AgamP3) reference genome obtained from VectorBase www.vectorbase.org was also included. Fontaine et al,. [10] made a whole genome alignment using ROAST [80] that represents approximately 40% of the euchromatic genome. We download the resulting MAF based on field-collected samples from http://datadryad.org/resource/doi:10.5061/dryad.f4114 [10]. We then added to this MAF our *An. fontenillei* assembly, and the highest coverage *An. bwambae* genome sequences available (see below).

***Anopheles fontenillei.*** We first generated a database with the scaffolds of the *An. fontenillei* assembly. Then we run blastn for each region in the MAF using AgamP3 as query sequence against the *An. fontenillei* scaffold database. We then repeat this blastn analysis using other species of the MAF regions as query; *An. arabiensis*, *An. quadriannulatus*, *An. melas* and *An. merus* (Text S2.1.1-Text S2.1.4). We did not repeat the analysis using *An. coluzzii* and *An. gambiae* as queries, due to its similarity to the reference genome AgamP3, or the two outgroup species, that are too divergent. We then selected the MAF regions for which we get one unique hit in any of the species, which represents the 63.2% of all the MAF regions for the eight species (Table S10). For the additional MAF region that gave more than one hit we exclude the multiple hits with *e-value* > 10^-4^ or with ≤ 40% of the query covered for each region in each species (Text S2.1.5) and, we recovered the sequences that became a unique hit after this filtering (Table S10). In total, we were able to include *An. fontenillei* in 75.2% of the previous MAF regions, which represent the ~30% of the euchromatic genome. For each of these MAF regions we cut the scaffolds according to the blast result information (Text S2.1.6). Then, we added these sequences to the corresponding MAF region using MAFFT as an aligner (v7.221, [81]). We use the function *-add* to modify as less as possible the initial MAF [82]. Finally, we joined each region of the MAF and generated the new MAF including the *An. fontenillei*genome (Text S2.2).

***Anopheles bwambae.*** We downloaded the tree individual sequences of *Anopheles bwambae* available at NCBI with fastq-dump; i) *An. bwambae 1*, SRR1255391, SRR1255392, and, SRR1255303, ii) *An. bwambae 3*, SRR1255390, and, iii) *An. bwambae 4*, SRR1255325. We then joined the SRR for individual 1 (Text S2.3). For each individual, we evaluated read quality with fastQC and trimmed the reads using cutadapt (v. 1.8.3; [83])(Text S2.4.1-Text S2.4.3). After the trimming, the quality per base was always higher than 24. We then map the trimmed read to AgamP3 reference genome using bwa-mem [84]. We performed several post-mapping steps including marking duplicates and realigning around indels using Picard (v. 1.109; http://picard.sourceforge.net), samtools (v. 1.3; [85]) and GATK (v3.4-46; [86])(Text S2.4.4-Text S2.4.8). Among the three available *An. bwambae* individuals we selected the one with the highest coverage, *An. bwambae* 1 with 33.2X, to add it to the MAF (the other two: *An. bwambae* 3, 11.7X and *An. bwambae* 4, 11.2X). We made a consensus sequence of the *An. bwambae 1* reads mapping to the AgamP3 sequence of every MAF regions with the 9 species (*An. gambiae*, *An. coluzzii*, *An. merus*, *An. melas*, *An. quadriannulatus*, *An. arabiensis*, *An. fontenillei*, *An. christyi* and *An. epiroticus*), using SAMtools mpileup (Text S2.5). If for a MAF region there were not *An. bwambae* reads, we added gaps so that we kept the same number of MAF regions as before. This had a marginal effect as it only occurred in 0.06% of all the MAF regions. Finally, we used MAFFT aligner, with the function *-add*, to add each consensus sequence of *An. bwambae 1* to the MAF regions and then joined all these regions in a new MAF (Text S2.2).

**MAF with four *An. fontenillei* individuals and three *An. bwambae* individuals.**To check the phylogenetic relationship between *An. fontenillie* and *An. bwambae,* we also created an additional MAF that included the 8 species previously available, the four *An. fontenillei* individuals, and the three *An. bwambae* individuals. We mapped each one of the seven individuals to AgamP3 reference genome as described previously for *An. bwambae* (Text S2.4). Then, we generated a consensus sequence for each of the new individuals for each of the MAF regions using SAMtools mpileup (Text S2.5). Finally, we add sequentially each of the new sequences to the available multiple alignment regions using MAFFT *-add* function as aligner (v7.221, [81, 82], Text S2.2). Finally, we joined all these information in a new MAF file.

**Window-based phylogenies.**We generated 50 kb genome-wide non-overlapping windows from the MAF (Text S2.6.1). For each window, we generated a maximum likelihood (ML) phylogenetic tree using RAxML (v8.2.4, [87]) with GTRGAMMA model and bootstrapping for 1,000 replicates (Text S2.6.2) [10]. We used the closer related species, *An. christyi*, as an outgroup because Fontaine et al. [10] already showed that the choice of the outgroup did not substantially alter the results. We excluded the windows with less than 10% of informative base pairs (e.g. < 5,000 bp) (following [10]). The different topologies obtained were sorted, counted, and analyzed using ad-hoc perl scripts (Text S2.6.3).

### Pairwise distance and bootstrapping

We used the R package ‘APE’ (v4.1, [88]) to estimate pairwise genetic distances based in the ML phylogenetic trees. We then performed the bootstrap analysis using the ‘boot’ package in R [89](Text S2.7).

### Centromeric regions alignment quality

For each chromosomal arm, we choose randomly 30 windows from centromeric regions and 30 regions from other genomic regions. Centromeric regions were defined based on the observed ancestry pattern in Fig. 3: 2L: 0 to 10Mb, 2R: 50 to 61.3Mb, 3L: 0 to 10Mb, 3R: 40 to 53.1Mb and X: 15 to 20.2Mb. For these 60 windows by chromosome arm, we gathered: i) the alignment length, ii) alignment length without completely undetermined characters and gaps, iii) proportion of gaps, and iv) the alignment patterns from the RAxML information file.

### Data analysis

We used R v3.2.5 (R Development Core Team, http://cran.r-project.org/) to perform all the statistical analysis. We used Inkscape software for figure edition (https://inkscape.org).

## Acknowledgements

We are grateful to P. Nosil and G. Lanzaro for comments on the manuscript. We thank the “Agence Nationale de la Preservation de la Nature” (ANPN), the “Station d’Etudes des Gorilles et Chimpanzes” (SEGC) and the “Centre National de la Recherche Scientifique et Technologique of Gabon” (CENAREST) that authorized this study and facilitated the access to the national parks of La Lopé. We specially thank Vincenzo Petrarca for its help interpreting chromosome polymorphism. Funding was provided by the “Institut de Recherche pour le Developpement”, the “Agence Universitaire de la Francophonie” (grant: OKANDA), the “Centre National de la Recherche Scientifique” (CNRS) and the “Consejo Superior de Investigaciones Cientificas” (CSIC) (grant PICS ANCESTRAL to DA and JG), the “ANR” (grant ANR-18-CE35-0002-01-WILDING to DA), and the “Ministerio de Ciencia, Innovation y Universidades/AEI” (grant BFU2017-82937-P to JG).

## Author contribution

**Conceptualization** DA, JG, CP, MGB

**Data Curation** MGB, DA, JG

**Formal Analysis** MGB, JG, DA,

**Funding Acquisition** JG, DA

**Investigation** PK, OAE, NR, PK, MFN, TWB, MP, DA

**Project Administration** DA, JG

**Resources** DA

**Supervision** JG, DA

**Validation** CC, FS

**Visualization** MGB, JG, NR, CP, DA

**Writing-Original Draft Preparation** MGB, JG, DA

**Writing-Review & Editing** MGB, JG, DA, CP

## Competing interests

The authors declare no competing interests.

**Data accessibility:**DNA sequences have been deposited to GenBank under accessions xxxxxx.

